# Gene validation and remodelling using proteogenomics of *Phytophthora cinnamomi*, the causal agent of Dieback

**DOI:** 10.1101/2020.10.25.354530

**Authors:** Christina E. Andronis, James K. Hane, Scott Bringans, Giles E. S. Hardy, Silke Jacques, Richard Lipscombe, Kar-Chun Tan

**Affiliations:** Centre for Crop and Disease Management, Curtin University, Bentley, WA, Australia; Curtin Institute for Computation, Faculty of Science and Engineering, Curtin University, Perth, WA, Australia; Proteomics International, Nedlands, WA, Australia; Centre for Phytophthora Science and Management, Murdoch University, Murdoch, WA, Australia

**Keywords:** Proteogenomics, oomycete, *Phytophthora*, Dieback, proteomics

## Abstract

*Phytophthora cinnamomi* is a pathogenic oomycete that causes plant dieback disease across a range of natural ecosystems and in many agriculturally important crops on a global scale. An annotated draft genome sequence and annotation is publicly available (JGI Mycocosm) and suggests 26,131 gene models. In this study, soluble mycelial, extracellular (secretome) and zoospore proteins of *P. cinnamomi* were exploited to refine the genome by correcting gene annotations and discovering novel genes. By implementing the diverse set of sub-proteomes into a generated proteogenomics pipeline, we were able to improve the *P. cinnamomi* genome. Liquid chromatography mass spectrometry was used to obtain high confidence peptides with spectral matching to both the annotated genome and a generated 6-frame translation. 2,764 annotations from the draft genome were confirmed by spectral matching. Using a proteogenomic pipeline, mass spectra were used to edit the *P. cinnamomi* genome and allowed identification of 23 new gene models and 60 edited gene features using high confidence peptides obtained by mass spectrometry, suggesting a rate of incorrect annotations of 3% of the detectable proteome. The novel features were further validated by total peptide support, alongside functional analysis including the use of Gene Ontology and functional domain identification. We demonstrated the use of spectral data in combination with our proteogenomics pipeline can be used to improve the genome of important plant diseases and identify biologically relevant missed genes. This study presents the first use of spectral data to edit and manually annotate an oomycete pathogen.

## Background

The primary role of a genome sequence is to elucidate the entire set of genes expressed by an organism. *In silico* prediction platforms are the main methods for predicting reliable gene sets. However, they can be problematic as transcriptome data does not always correlate with the protein products and their abundance [1]. Curating genes correctly and accurately is fundamental in defining the biochemical composition of an organism [2]. Sequence transcripts and orthologues from related and similar organisms are the primary methods in accurately predicting such genes and identifying interesting and relevant biological components [3]. Evidence-based curation includes transcript data and associated functional annotation such as Gene Ontology (GO) and Protein Families (PFAM) based on sequence homology to other related species [4, 5]. The challenges in defining comprehensive gene products result in under-represented annotations and incorrectly defined exon boundaries that can miss biologically important features of a genome.

Proteogenomics is a proven but underutilised technology that integrates high confidence peptide data derived from mass spectrometry analysis with genomics as a method to improve gene annotation [6–8]. Proteogenomic pipelines have been used in phytopathogenic fungi such as *Parastagonospora nodorum*, where mass spectra were used to validate transcriptomic data, edit the genome annotation and identify new candidate genes, generating a more accurate genome that can be used for downstream work [9, 10]. Proteogenomic analysis has also allowed the identification of potential effector molecules in fungi, which has important implications for characterising virulence and understanding the plant-host interface. The proteome of the causal agent of black spot in pear, *Venturia pirina,* was analysed by mass spectrometry and 1085 novel protein groups were identified, 14 of which were fungal candidate effector genes [11]. This provides useful insight into the mechanisms of pathogenicity and has the potential to be exploited to control oomycete and fungal plant pathogens.

Mass spectrometry based proteomics can also be used to overcome some of the constraints of traditional gene prediction methods and with continuing advances in proteomics technologies becomes a desirable tool to elucidate the biochemistry of an organism [12]. High throughput proteomic pipelines such as liquid chromatography mass spectrometry (LC-MS/MS) are becoming more sensitive, rapid and less expensive [13]. As an added benefit to transcriptome work, quantitative proteomics can inform on differential expression of proteins [14].

*Phytophthora cinnamomi* is a phytopathogenic oomycete that causes dieback and root rot in natural and agricultural systems across the globe. Its hosts include many species of native Australian flora, as well as crops such as avocado and macadamia [15]. Oomycetes proliferate by releasing motile, asexual units of reproduction called zoospores. When temperatures and humidity reach favourable levels, *P. cinnamomi* produces fruiting bodies called sporangia which expel a free swimming zoospores in to the environment, allowing the organism to spread between susceptible hosts. As the zoospores colonise a hosts root system, mature structures including those required for sexual reproduction and nutrient acquisition form, eventually killing the host. Due to the host range, ability to survive harsh environmental conditions and aggressive pathogenicity, *Phytophthora* is recognised as one of the most economically important oomycete genera, with insufficient existing control strategies to minimize its impacts [15, 16].

*P. cinnamomi* is controlled by the application of phosphite to vegetation [17]. The mechanisms involved in phosphite are not well understood, however it is suggested that it has both direct influence on the organism and indirect action through the plant by priming host defence responses [18–20]. Genes involving cell wall synthesis and cytoskeleton function have been suggested to be disrupted in *P. cinnamomi* when treated with phosphite *in vitro* [21]. Despite its economic and ecological importance, little is known about the molecular mechanism of *P. cinnamomi* phytopathogenicity. It is hypothesised that *P. cinnamomi* secretes effectors based on studies on other oomycetes such as *P. infestans* and *P. ramorum* [22]. Virulence and infection related molecules such as β-cinnamomin have been identified in *P. cinnamomi* [23].

A genome sequence of the West Australian *Phytophthora cinnamomi* MU94-48 isolate (GenBank accession JX113294) was established (unpublished and publicly available at https://mycocosm.jgi.doe.gov/Phyci1) [24]. This is a valuable tool that can be used to identify effectors and elucidate the molecular mechanisms of virulence [25]. The version 1 (V1) assembly has a coverage of 69.6x and comprises 9,537 contigs, 1,314 scaffolds with 26,131 predicted gene models. The predicted gene models of *P. cinnamomi* are inflated compared to many *Phytophthora* species such as *P. infestans, P. ramorum, P.capsici* and the more closely related *P. sojae,* which have reported 17,797, 16,066, 19,805 and 15,743 gene models respectively [25].

Proteomics data has proven a useful tool to improve the genome annotation of several phytopathogens where high quality mass spectra complemented transcriptomic data and identified potential annotation inaccuracies of exon boundaries and unsuspected gene models. We aimed to use spectral data from several *P. cinnamomi* sub-proteomes to assist in gene calling. These sub-proteomes represent a wide coverage of the *P. cinnamomi* proteome and include a diverse repertoire of soluble proteins. Zoospores characterize the infective life stage and the extracellular proteome is likely to contain proteins related to virulence. These were analysed by 2D LC-MS/MS and resulting spectra were matched to the current gene prediction models. To generate a list of peptides which potentially do not match current models, a 6-frame translation was generated and used for spectral matching. A list of peptides indicating potential altered or novel gene models was generated using the genomic coordinates and peptide open reading frames. These were subsequently used to carefully manually edit current annotations and curate novel features on a homology basis with proteins of similar species. Using this proteomics dataset, we refined the genome for downstream proteomic work which will aid the identification of virulence factors and metabolic targets for chemical control. By working towards completing the *P. cinnamomi* genome, downstream proteomic work will be more accurate as the gene set is more representative of what is being expressed. There is also the potential for effector virulence gene discovery and improved biochemical characterisation which can lead to development of resistance gene inclusion in hosts and more targeted methods of chemical control.

## Methods

### Growth and maintenance of *P. cinnamomi*

*P. cinnamomi* MU94-48 (Centre for Phytophthora Science and Management, Murdoch University, Western Australia) stocks plugs were stored in sterile water in McCartney bottles and grown on V8 agar at room temperature in the dark. For mycelial, secretome and zoospore production, four plates were used for each biological replicate. Four days of mycelial growth were scraped from the plate and were inoculated into Riberio’s minimal media supplemented with 25 mM glucose [26]. The cells were incubated for 3 days at 24°C in the dark. Mycelia were isolated by centrifugation and the culture filtrate containing secreted proteins was decanted and filter sterilised. The mycelial pellet was washed twice with MilliQ water and observed microscopically to confirm that hyphal cells predominated. Zoospores were produced as previously described by Byrt & Grant, isolated from mycelial fragments by passage through a glass wool syringe, observed microscopically to ensure purity, and counted with a haemocytometer [27]. Approximately 4.8E5 spores were used for each biological replicate.

### Protein extraction

Mycelia and zoospores were ground using mortar and pestle in liquid nitrogen and extraction buffer used to extract and solubilise proteins as previously described [28]. Samples were kept on ice for 30 minutes with regular gentle mixing and centrifuged at 20,000g at 4°C for 30 minutes. The protein solutions were subsequently desalted and protein amount was estimated using Direct Detect cards (Merck Millipore, Darmstadt). All samples were freeze dried before further processing. SDS-PAGE was performed for all samples to ensure proteolysis was minimal.

### Sample preparation

To visualise each sub-proteome, 20 μg of each sample was loaded onto a 1D SDS-PAGE. To determine the amount of intracellular contamination in the extracellular proteome, the activity of an intracellular enzyme marker glyceraldehyde phosphate dehydrogenase (GAPDH) was assayed on each sub-proteome as per the manufacturer’s instructions (Sigma, St Louis). 500 ug of each sample was resuspended in 250 uL 0.5 M triethylammonium bicarbonate (pH 8.5) before reduction and alkylation with 25 uL of 50 mM tris(2-carboxyethyl)phosphine (Thermo Scientific, Waltham) and 12.5 uL 200 mM methyl methanethiosulfonate (Sigma, St Louis) respectively. Samples were digested overnight at 37°C with trypsin (Sigma, St Louis) at a ratio of 1:10, subsequently desalted on a Strata-X 33 um polymeric reverse phase column (Phenomenex, Torrance, CA, USA) and dried in a vacuum centrifuge.

### High pH reverse phase chromatography

Dried peptides were separated by high pH reverse phased liquid chromatography on an Agilent 1100 HPLC system using a Zorbax Eclipse column (2.1 × 150 mm, 5 um) (Agilent Technologies, Palo Alto). Peptides were eluted with a linear gradient of 20 mM ammonium formate pH 10, 90 % acetonitrile over 80 minutes. A total of 98 fractions were collected, concatenated into 12 fractions based on collection order and dried in a vacuum centrifuge. The UV trace was also used to visualise the total peptide content and depth of each sub-proteome.

### Nano LCMS/MS

Fractions were resuspended in 100 uL of 2% acetonitrile and 0.1% formic acid and loaded onto a Shimadzu Prominence nano HPLC system (Shimadzu, Kyoto, Japan). Peptides were resolved with a gradient of 10-40% acetonitrile (0.1% formic acid) at 300 nL/min over 180 minutes and eluted through a nanospray interface into a 5600 TripleTOF mass spectrometer (AB Sciex, Framingham, MA). The data was acquired in an information-dependent acquisition mode with Analyst TF 1.6 software (AB Sciex, Framingham, MA). The MS settings were as follows: Ionspray Voltage Floating = 2300 V, curtain gas = 20, ion source gas 1 = 20, interface heater temperature = 150, and declustering potential = 70 V. The TOF MS scan was performed in the mass range of 400-1250 Da with a 0.25 s TOF MS accumulation time, whereas the MS/MS product ion scan was performed in the mass range of 100 - 1800 Da with a 0.1 s accumulation time. The criteria for product ion fragmentation were set as follows: ions (>400 *m/z* and <1250 *m/z*) with charge states of between 2 and 5 and an abundance threshold of >250 cps. Former target ions were excluded for 10 s after one occurrence. The maximum number of candidate ions per cycle was 20.

### Data analysis

Mass spectral data were analysed using Protein Pilot 4.5 Beta Software (July 2012; Sciex). MS/MS spectra were searched against the genomic proteins and the 6-frame translated data set constructed from the genomic assembly scaffolds using EMBOSS: getorf (v6.6). Search parameters were: Cys Alkylation: MMTS; Digestion: Trypsin; Instrument: TripleTOF 5600; Special factors: None; Quantitation tab checked: Bias correction and Background correction tabs checked; ID focus: Biological modifications; Search effort: Thorough; Detected protein threshold (Unused ProtScore (CONF)): 0.05 (10%); False discovery rate analysis tab checked. All identified proteins had an Unused Protscore of >1.3 (peptides identified with >95% confidence), as calculated by the software and a global false discovery rate of <0.1% determined at the protein level. To determine the sub-proteome enrichment, the resulting sequences of matched proteins were analysed using the protein localisation tool WolfPSORT (version 0.2, plant parameters) [29]. Proteins were assigned to a predicted sub-cellular location based on sorting signals, amino acid composition and functional motifs.

### *De novo* proteogenomics

Peptide matches to the 6-frame translated assembly were mapped back to their genomic location and a set of criteria described below were applied to determine which genes suggest incorrect boundary annotations and which peptides support discovery of new genes. Firstly, BEDtools (version 2.28.0, 2019) was used to distinguish peptides into the following groups using the intersect and subtract features: a) peptides more than 200 base pairs from coding regions of genes (CDS), b) peptides within 200 base pairs from CDS features but do not overlap CDS boundaries, c) Peptides that overlap CDS boundaries, and d) peptides that remain within CDS feature boundaries [9, 30]. Subsequently, the CDS Mapper tool (version 0.6, 2011, https://sourceforge.net/projects/cdsmapper/) was used with default parameters to further classify these based on their frame match to corresponding CDS features of the annotated draft genome (Figure 2) (Bringans et al., 2009). All peptides suggesting novel or altered gene models were blasted (BLASTp, version 2.9.0, 2019) with the following search parameters: organism: *Phytophthora* (tax ID 4783), expect threshold: 2E5, word size: 2, matrix: PAM30, gap costs: existence 9 and extension 1. All peptides with significant returns (e<1E-3) were considered for manual annotation. Peptides that did not return significant results were not used for this analysis. All significant BLAST hits were transferred onto the *Phytophthora* draft genome and manually edited to comply with sequence features such as start/stop codons and non-sequenced regions. The novel annotated genes were further analysed for total number of supporting peptides (as per Protein Pilot methods described above). Genes that had only one high confidence peptide were included for the purposes of gene discovery [31]. Protein Family domains (PFAM), Gene Ontology (GO) terms and Kyoto Encyclopedia of Genes and Genomes (KEGG) were also assigned using Interpro scan (version 5.44-79.0, 2020) using and EGGNOG-mapper (version 2, 2019) using default parameters. To determine whether any pathogenesis related proteins were present within the novel set, annotations were analysed for presence of any potential virulence factors using PHI-BASE (version 4.9, 2020) using default parameters [32]. The Codon Adaptation Indexes (CAIs) of each novel gene were also calculated using Emboss CAI (version 6.6, default parameters), which indicated gene annotation with anomalous usage of codons [33].

## Results

### Sub-proteome enrichment

To obtain a representative proteome of *P. cinnamomi*, vegetative mycelia and transient short-lived zoospores of *P. cinnamomi* were used as these are the dominant cell types that grow and initiate infection in hosts. In addition, we extracted soluble secreted proteins (secretome) from the mycelia, which are widely studied due to their implications on pathogen-host interactions. The purity of the mycelia and zoospores was observed under a stereoscope (Figure 1). Figure 1A shows no evidence of intercellular contamination and demonstrated the purity of these cell types. The large mass of mycelia had not produced zoospores or their precursor (the sporangia) in this method of *in vitro* cell culture. Similarly, vegetative mycelia was not observed in the zoospore preparation (Figure 1B).

**Figure 1.**
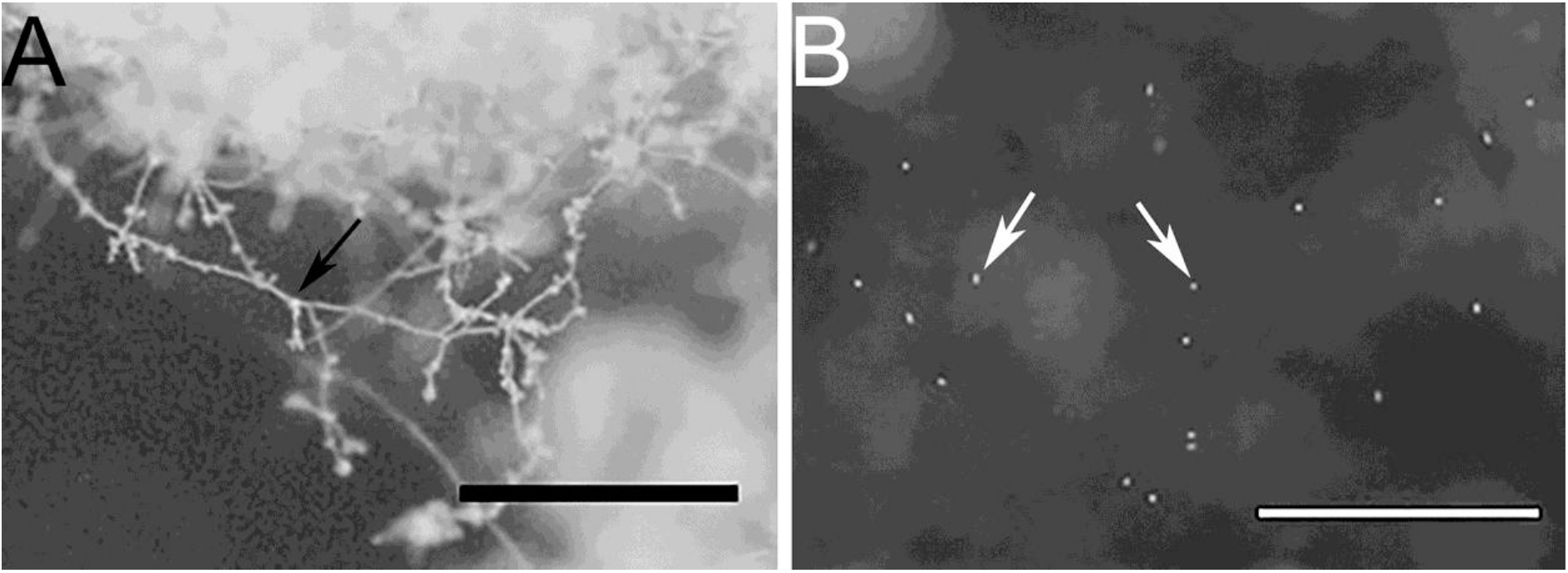
Stereo microscope images of A) a mycelial mass, and B) free swimming zoospores, indicating minimal to no cross contamination between cell types. Bars represent 1 mm scale. Black arrow indicates mycelia and white arrows indicate zoospores.

1D SDS-PAGE was run to visualize the sub-proteomes of each cell type (Figure 2A). The banding patterns of each sub-proteome show differences in total protein content. The extracellular proteome showed enrichment in lower molecular weight proteins whereas the mycelia and zoospores had proteins that spanned over the whole mass range. To test the purity of the secretome, an enzyme activity assay of the cytoplasmic marker GAPDH was measured, which should only be present in small amounts (Figure 3b) [34]. Both the mycelia and zoospores had similar detected amounts of GAPDH detected, approximately 4.7 and 4.8 mU/mg protein, respectively. GAPDH was also detected in the secretome, however at lower amounts (1.6 mU/mg protein). The RP-HPLC UV total ion count traces indicated differing protein content between the three sub-proteomes, as majority of the peaks do not match in intensity and retention time (Figure 3c). The majority of proteins detected in the mycelial and zoospore were localised intracellularly at 45% and 41%, respectively as predicted by WolfPSORT (Table 1). The secretome was enriched in extracellular localisation proteins with a predicted 18% compared to 5% in both the mycelia and zoospores.

**Figure 2.**
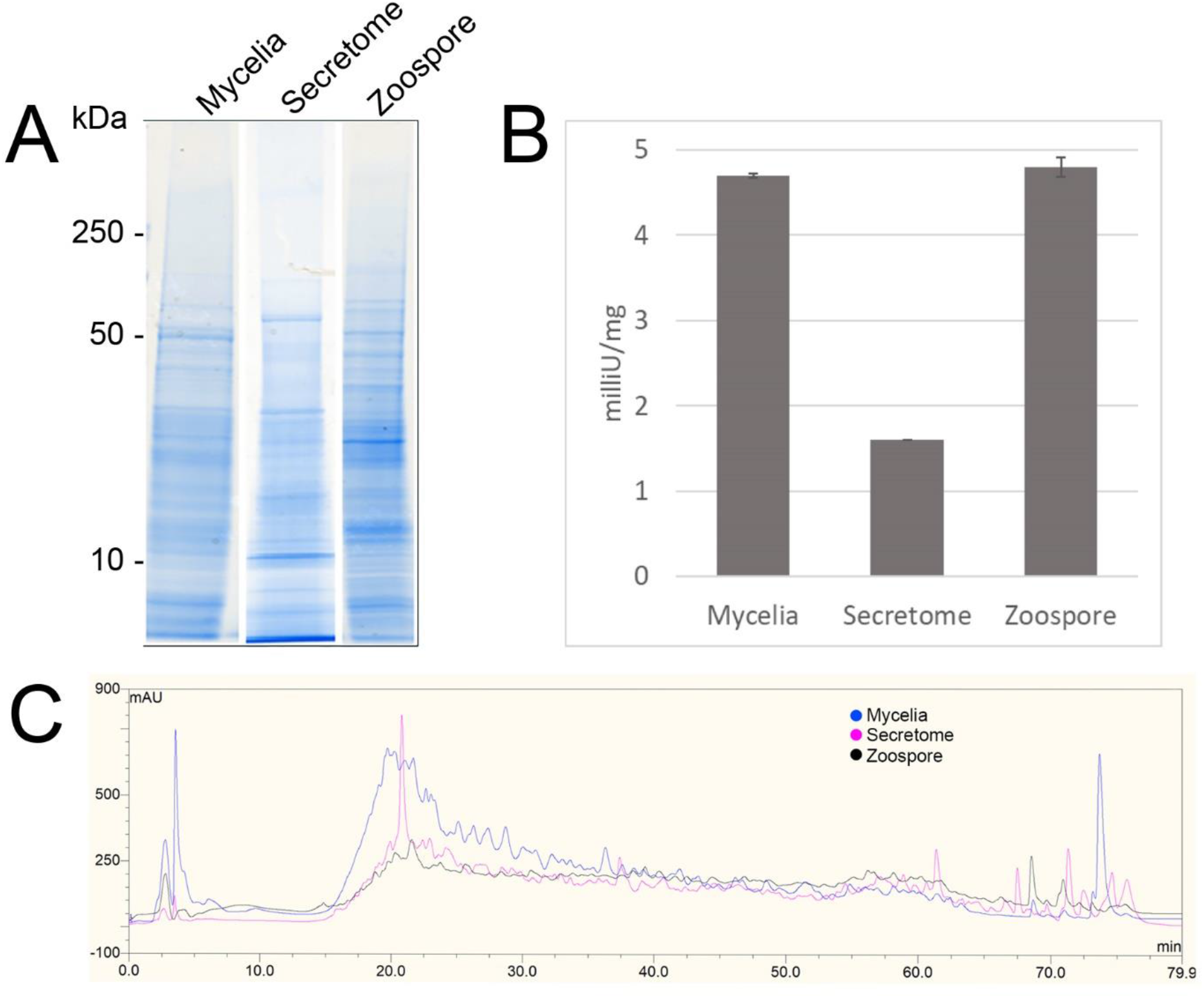
Quality control of sub-proteomes. A) SDS-PAGE analysis of *P. cinnamomi* sub-proteomes. B) Glyceraldehyde phosphate dehydrogenase activity within each sub-proteome indicating relative levels of contamination of intracellular proteins in the extracellular proteome. C) High pH reverse phase HPLC separation of 500μg of each sub-proteome demonstrating sufficient peptide separation and differing protein content.

**Figure 3.**
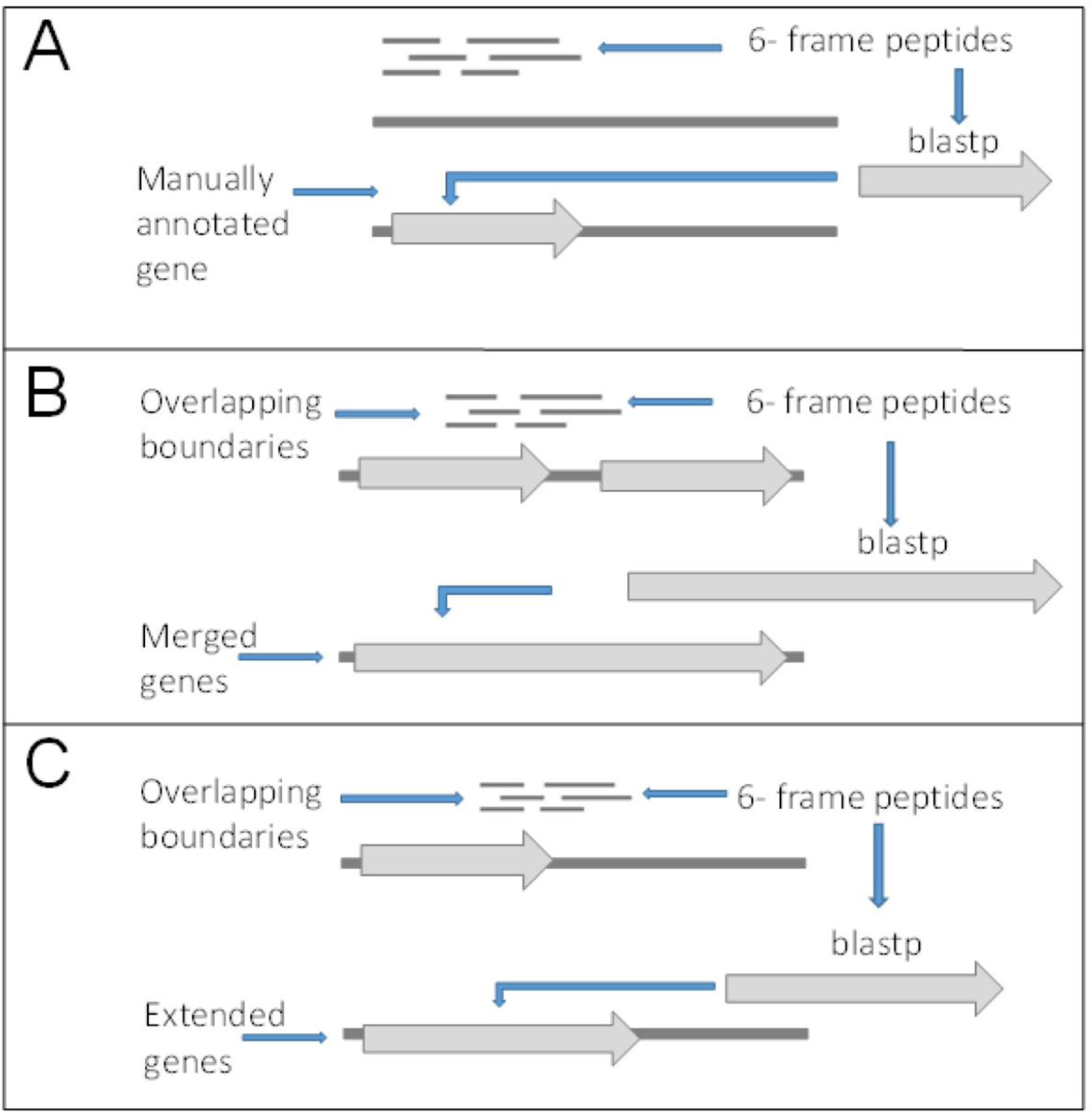
Criteria for gene editing and discovery. A) Peptides matching to 6-frame open reading frames in loci with no surrounding genes or out of frame from surrounding genes and returned significant BLASTp hits were manually annotated as new genes. B) Peptides overlapping boundaries of multiple genes within the same frame and returned significant BLASTp hits were used to manually merged genes. C) Genes with in frame peptides overlapping boundaries and returned significant BLASTp hits were manually edited.

**Table 1.** WolfPSORT localisation prediction of the whole genome annotation and sub-proteomes.

| WolfP SORT | Total no. proteins | Predicted Intracellular | Predicted Extracellular | Low confidence prediction |
| --- | --- | --- | --- | --- |
| Whole genome prediction | 26,131 | 43 % | 6 % | 51 % |
| Mycelial proteome | 3,209 | 46 % | 5 % | 49 % |
| Secretome proteome | 1,605 | 42 % | 18 % | 40 % |
| Zoospore proteome | 2,304 | 50 % | 5 % | 45 % |

### Validation of V1 gene models using sub-proteome spectra

The mass spectra were used to validate the draft annotation of the *P. cinnamomi* genome. The annotations acquired from JGI Mycocosm (assembly annotation version 1.0) were designated in this study as ‘V1’ and the annotation set containing subsequently manually edited loci was designated ‘V2’.

Non-redundant peptide matches (at least two 95% confident peptides) resulted in 2,554, 1,362, and 2,304 proteins from the mycelia, secretome and zoospores respectively. From this data, 2,764 unique proteins from the V1 predicted gene set were identified (Figure 4). 526, 215 and 432 proteins were unique to the mycelia, secretome and zoospores respectively, which implies a wide range of the whole proteome detected. The mycelia and zoospores had more unique protein identifications than the secretome, which may be a result of an expected lower mass range of an extracellular proteome that were below the acquisition detection limits.

**Figure 4.**
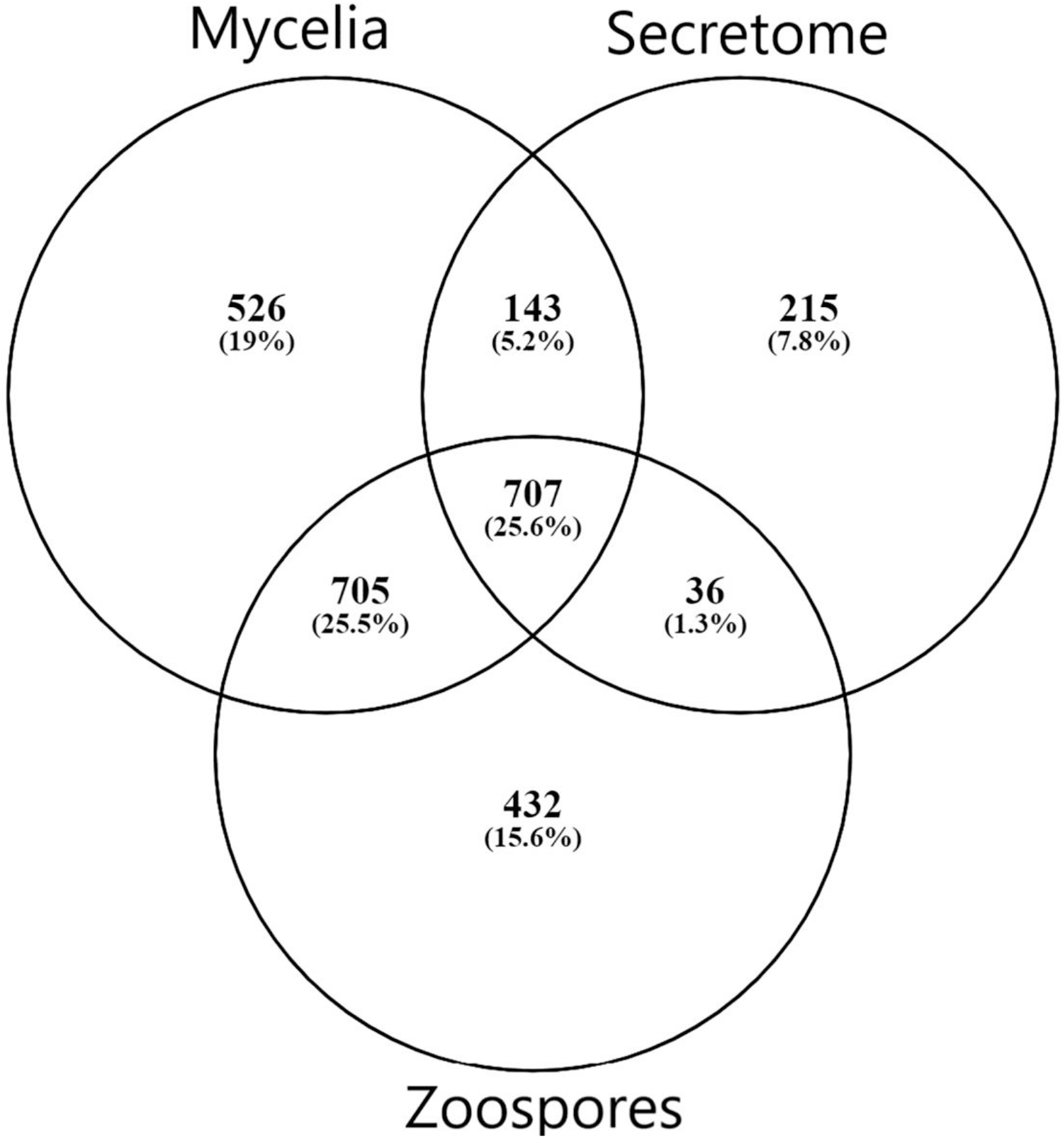
Number of proteins detected by mass spectrometry of each sub-proteome supported by at least two 95% confident peptides.

**Figure 5.**
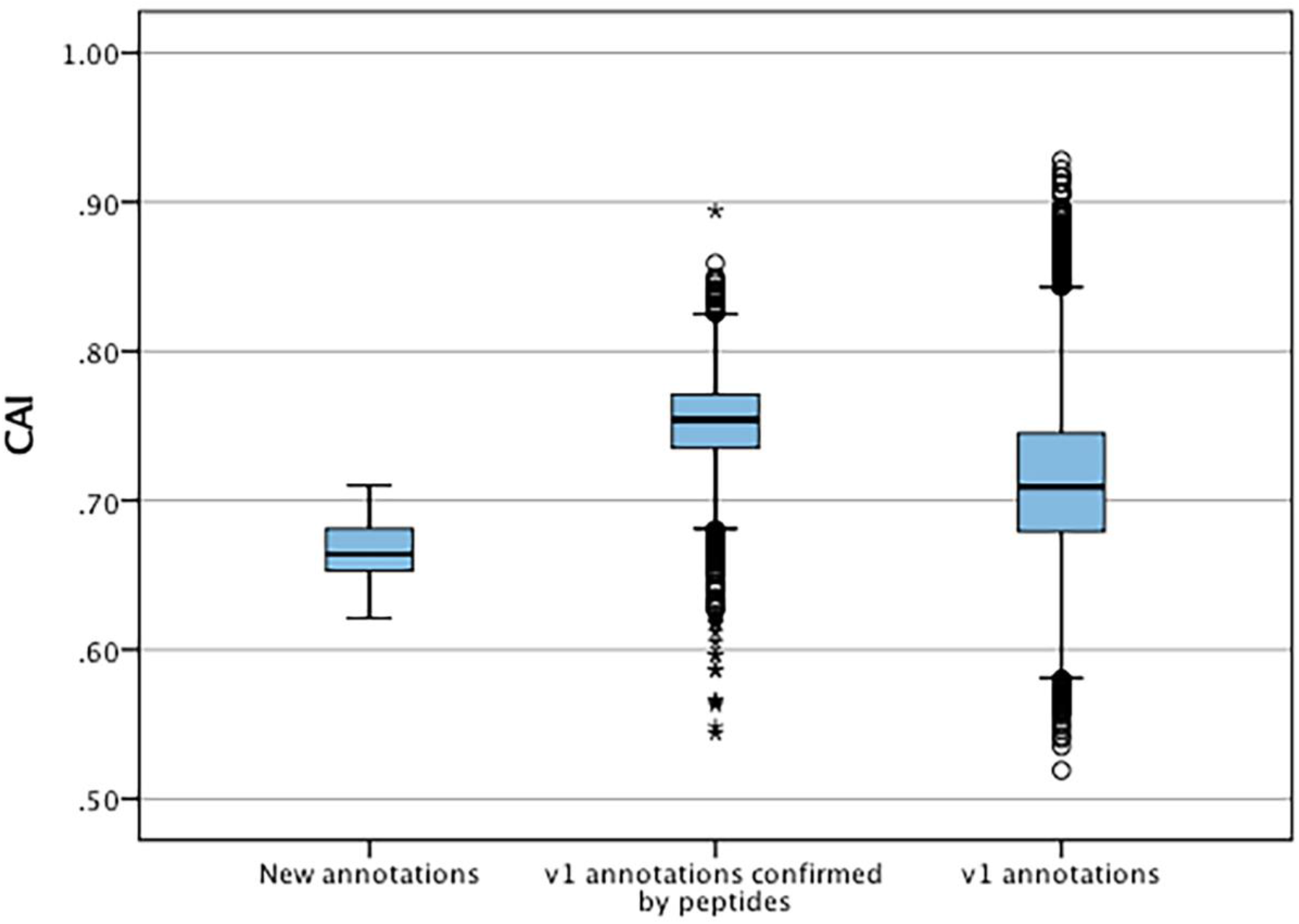
Distribution of codon adaptation indices between new annotation and v1 annotations confirmed by mass spectrometry and all v1 annotations. * represent significant differences (p <0.05) of data sets to V1 annotations.

When matched to 4,874,027 generated open reading frames (ORF) of the 6-frame translation, 2,752, 1,355, and 2,334 ORFs from the mycelia, secretome and zoospores were identified (Table 2). Although this does allow us to match more peptides to the genome than the V1 annotation, some level of redundancy is expected from matching to reading frames that do not form genes. The false discovery rate for all mass spectra analysis was <0.1% using the Protein Pilot decoy database method, which is within the limits of the general consensus for large scale proteomic data [35, 36]. Of the V1 detected by mass spectrometry, 2,398 had additional support by assigned GO terms and/or PFAM domain.

**Table 2.**
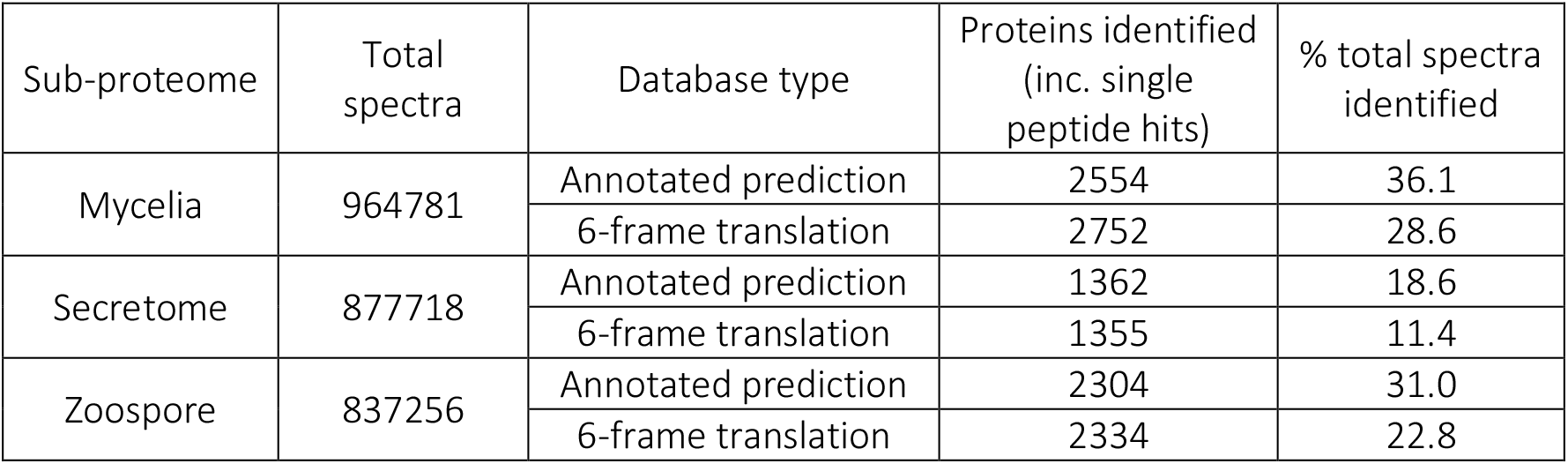
Summary of mass spectra identification using the annotated protein and 6-frame open reading frame databases.

| Sub-proteome | Total spectra | Database type | Proteins identified (inc. single peptide hits) | % total spectra identified |
| --- | --- | --- | --- | --- |
| Mycelia | 964781 | Annotated prediction | 2554 | 36.1 |
|  |  | 6-frame translation | 2752 | 28.6 |
| Secretome | 877718 | Annotated prediction | 1362 | 18.6 |
|  |  | 6-frame translation | 1355 | 11.4 |
| Zoospore | 837256 | Annotated prediction | 2304 | 31.0 |
|  |  | 6-frame translation | 2334 | 22.8 |

**Table 3.** Confirmation of genes supported by peptides within or crossing exon boundaries.

|  |  |
| --- | --- |
| **Total number of genes** | **26478** |
| Genes confirmed by spectral matching (Protein Pilot) | 3468 |
| Genes with 6-frame peptide support- with boundary conflicts | 52 |
| Genes with no 6-frame peptide support (inc. with boundary conflicts) | 19724 |
| Genes with only mismatched 6-frame peptides | 69 |
| Genes with matched and mismatched 6-frame peptides | 24 |

### Annotating new gene models by homology criteria

Although there is peptide support for a large number of the V1 genes, it is expected that there are some forms of incorrect intron and exon boundary annotations that can be detected using spectral data. In addition, this spectral data can also be used in the detection of new genes. 23,457 unique high confidence peptides matched to the 6-frame ORFs were mapped back to their genomic location. 22,443 peptides mapped completely within coding exon boundaries. 274 peptides mapped partly within exons (i.e span across boundaries) and 287 within 200 bp of boundaries (Figure 2B, 2C). 453 peptides mapped more than 200 bp from exon boundaries (Figure 2A). Furthermore, the frame test applied more stringent criteria for frame matching of these peptides to corresponding V1 annotations (Table 4). A total of 1,010 peptides did not match the frame of corresponding CDS features or were further than 200 bp from any gene models. This suggested 438 gene features with potentially incorrect boundaries. These were considered as candidates for new gene models.

**Table 4.** Summary of frame matches of peptides with nearby CDS features and number of significant Blastp hits (E value > 1E-3) and number of gene features manually

|  | Frame match (CDS) | Frame mismatch (CDS) | Total peptides | No. significant BLAST hits of conflicting peptides |
| --- | --- | --- | --- | --- |
| Peptides within CDS boundaries | 22134 (3178) | 309 (242) | 22443 | 38 |
| Peptides crossing CDS boundaries | 240 (171) | 34 (31) | 274 | 129 |
| Peptides within 200bp from CDS boundaries | 73 (52) | 214 (165) | 287 | 67 |
| Peptides more than 200bp from CDS features | - | - | 453 | 118 |

To select peptide candidates that would likely result in alteration of V1 genes and curation of new genes, Blastp was used. Peptides that returned significant hits to other *Phytophthora* species were used to manually edit and curate new genes (Table 4). This largely reduced the number of potential edited and new genes due to both the redundancies of 6-frame peptides and rigorous Blastp parameters used for peptide matches. Of those with conflicting boundaries, 70 peptides showed significant homology to other *Phytophthora* species. Of the peptides that were further than 200 bp from any gene, a total of 118 peptides returned significant BLASTp hits, suggesting the presence of previously unannotated genes on the *P. cinnamomi* genome. The homologous sequences were transferred onto the *P. cinnamomi* genome and the annotations were manually integrated, taking into consideration differences in the genome and features such as introns.

Using these criteria, a total of 60 genes were edited, which equates to an error rate of approximately 2% of the detected proteome. The CDS coordinates of the edited genes are shown in Additional file 1. Of these, 44 were modified by extending the exon boundaries and there were 16 instances of merged genes. Additionally, 23 new previously undefined genes were annotated (Table 5). These annotations were uploaded to the GenBank under accessions MT820663-MT820655. The edited annotations will be referred to by original annotation identification with ‘V2’ suffixed, as listed in the Additional files 2 and 3, respectively. In summary, we identified errors in 60 V1 genes which were manually altered and added a further 23 annotations to the gene set of *P. cinnamomi*.

**Table 5.** Summary of original predicted and newly annotated/ edited genes using proteogenomics

|  | No. genes |
| --- | --- |
| Version 1 prediction | 26131 |
| Total modified genes | 60 |
| Modified by extension | 44 |
| Modified by merging genes | 16 |
| New gene annotations | 23 |
| Total number of version 2 genes | 26151 |

### Validating edited and new genes

The edited genes were subsequently analysed for total peptide support and differences in functional assignment compared to the original annotation. Peptides within the edited regions were manually counted (Table 6). Of the extended genes, only one had no other supporting information other than the support of one 95% confident peptide in the extended portion of the gene (e_gw1.28.366.1_V2). All other extended genes had support from more than two high confidence peptides and/or homologous functional assignment. Similarly, only one merged gene had a single peptide supporting the merged region of the annotation (gw1.160.19.1_V2). All others were supported by two or more high confidence peptides, which is the general requirement for protein identification in proteomics [37]. Genes were analysed for GO terms, PFAM domains and KEGG orthologues (KO) to determine whether the altered boundaries change their functional annotation assignment (Table 6). Details of each functional annotation are shown in Additional files 2 and 3.

**Table 6.** Summary of new gene validation using supporting peptides, PFAM and Gene Ontology terms using Protein Pilot.

|  | Merged annotations | Extended annotations |
| --- | --- | --- |
| **Supported by only one peptide** | 6 | 8 |
| **Supported by two or more peptides** | 10 | 36 |
| Genes with PFAM domains | 13 | 39 |
| Edited genes with differing PFAM to v1 | 3 | 3 |
| Genes with GO terms | 13 | 35 |
| Edited genes with differing GO to v1 | 1 | 1 |
| Genes with KO assignment | 13 | 34 |
| v2 genes with differing KO to v1 | 0 | 1 |

The original mass spectra were matched to the set of new genes (using Protein Pilot- see methods) to determine how many peptides supported each gene (i.e. determine if any genes were a product of single peptide matches) (Table 7). Of the 23 new genes identified, one new gene had support from only one high confidence peptide (MT820633). All new genes were detected in the mycelia and most were also identified in the secretome and zoospore (Additional file 4). The remaining 22 genes had at least two or more supporting peptides.

**Table 7.** Summary of new gene validation using supporting peptides, Protein Families and Gene Ontology terms. Details of annotations of each entry are shown in Supplemental data 1

|  | Number of new gene models |
| --- | --- |
| **Supported by only one peptide** | **1** |
| **Supported by two or more peptides** | **22** |
| Contain PFAM domains | 21 |
| Contain GO terms | 21 |
| Containing KO | 17 |
| No functional support | 2 |

To further support this new gene set, protein sequences were analysed for protein function by assignment of PFAM domains, GO terms and KO assignment (Table 7). Details of these annotations for each entry are shown in Additional file 4. The new annotations were analysed for virulence factors using PHI-BASE. None of these annotations returned a significant hit to any known virulence factors.

### Codon adaptation Index

The codon adaptation indices were calculated for the set of new features and compared to the V1 gene set to identify significant differences in codon usage and distribution that could indicate possible causes for errors and missed genes (Figure 8). The distribution of the CAIs of the new set were significantly different (t-test, p value <0.05) than those of the predicted gene set suggesting a higher proportion of less common codon usage in the new set. These were also significantly lower to the CAIs of all original annotations that had high confidence supporting peptides. Each new gene was also analysed for unusual codon usage, primarily the use of start codons other than methionine and not terminated by a stop codon (Table 8). Only one new annotation MT820649 had abnormal codon usage, where there was no annotated start codon at the correct locus.

**Table 8.** Codon usage summary of the new gene set.

|  | No. features |
| --- | --- |
| Total features | 23 |
| Normal codon usage | 22 |
| Stop codon only (no start codon) | 1 |

## Discussion

The three sub-proteomes (mycelia, secretome and zoospores) represent a diverse range of proteins and capture the majority of the *P. cinnamomi* proteome. Although microscopic observation indicated successful purity in the mycelia and zoospores, the GAPDH assay showed some cytoplasmic contamination in the secretome. Cytoplasmic markers such as GAPDH and malate dehydrogenase have been used in other studies as indicators of intracellular contamination in secretome samples of other organisms [38, 39]. In these cases, some level of contamination was similarly observed in isolated secretome samples, likely due to some level of cellular leakage. Other known cytoplasmic markers such as malate dehydrogenase have been observed in fungal secretomes, where their extracellular functions are not known [40]. This set of enrichment data confirms that the sub-proteomes were sufficiently enriched for this study and the total proteome is diverse and represents both growing and infective stages of development.

The aim of using these three sub-proteomes was firstly to validate as much as the *P. cinnamomi* draft gene set as possible. Through spectral matching, we verified 10.6% of the predicted gene set. The differing sub-proteomes, as shown above, are reflected in the validation of V1 genes by spectral matching. Unique protein identifications of approximately 19%, 8% and 16% of the mycelia, secretome and zoospores respectively, accounted for the differences in observed SDS-PAGE banding patterns and RP-HPLC traces. Proteomic studies of other *Phytophthora* species indicated variable numbers of unique proteins to these sub-proteomes. A 2- dimensional proteomic study of the oomycete *P. palmivora* indicated 1% unique proteins for mycelia and zoospores [41]. However, a profile of the *P. infestans* secretome indicated similar coverage of extracellular proteins to this study [42].

Using the mass spectra matched to the 6-frame translation of the draft genome, we refined the draft genome of *P. cinnamomi*. Total peptide support and functional assignment were used to validate and to obtain the most accurate representation of edited and newly curated genes. We compared the functional assignments of V1 and V2 edited genes to identify differences inferred by the changes in annotation features. The PFAM domains associated with the extended genes that differed from V1 genes were involved in energy production and one recombination protein (gw1.193.42.1_V2 and fgenesh1_kg.277_#_5_V2). Similarly, the only differing GO was that of DNA repair (CE70043_1777_V2). The KO of V2 extended genes remained mostly the same with minor changes and one enzyme code, an enzyme involved in carbohydrate metabolism was not present in V1 genes. These changes include a PAMP recognition signalling factor in gw1.44.72.1_V2. The majority of merged gene features were mostly between CDS features of the same gene. Therefore, there were minimal functional assignment differences between V1 genes and those altered by merging. However, three instances merged whole genes (fgenesh1_kg.79_#_14_#_15_V2, estExt_fgenesh1_pm.C_90019, fgenesh1_pm.9_#_20, e_gw1.9.526.1_V2 and gw1.243.65.1, gw1.243.79.1_V2). fgenesh1_kg.79_#_14_#_15_V2 combined two whole genes from the V1 annotation, the V2 functional assignment included an additional PFAM domain, PF12698, an ABC-2 family transporter protein, which are often highly expressed in plant pathogens such as the oomycetes as they play roles in the biotrophic phase of infection and pathogenicity [43, 44]. The second, estExt_fgenesh1_pm.C_90019, fgenesh1_pm.9_#_20, e_gw1.9.526.1_V2, merged three whole genes, and included three different PFAM domains, two Poly(ADP-ribose polymerase domains and one WGR domain. This edit also resulted in one gene ontology difference, the presence of an NAD+ ADP-ribosyltransferase. There were no other differences in the GO and KEGG ontologies between V1 and V2 merged genes. Although these functional differences do not indicate major functional differences, they can impact the way in which we classify these proteins when trying to understand their role in a system.

The newly curated genes were validated using total peptide support, functional assignment and also examined for their codon usage to gain a better understanding of why they were missed in the V1 annotation. Although MT820633 had only one supporting peptide, it had PFAM, GO and KEGG assignment, all indicating its function to be associated with ankyrin, a protein family that is involved in the formation of the cytoskeleton and has been associated with signal transduction in other oomycete pathogens [45]. MT820636, MT820637 and MT820651 although had significant blast hits to other oomycetes but did not have any PFAM, GO or KO assignments. Functional domains were identified for all other new genes, most of which were related to general biochemical processes, including energy production, translation and transporter activity. PFAM domains and GO assignments that were associated with new genes but not present in V1 genes were mostly domains of ribosomal proteins and one ferredoxin type domain. Ribosomal proteins are highly conserved between species of oomycetes. This has been shown in *Pythium insidiosum* using expressed sequence tags, that show homology between several of the *Phytophthora* species [46, 47]. Ferredoxin domains have been identified in *P. parasitica* and were found to be associated with ATP generation [48]. Of the KO assignments, nine from V2 genes were not present in V1 genes. These were mostly associated with general metabolic and cellular functions, translation and genetic information processing functions, with many domains associated with ribosomes (Additional file 1).

PHI-BASE indicated no significant homology to virulence factors in the new gene set. This was expected as all cell culture was performed *in vitro* where there is minimal stimulation to produce and release virulence factors such as effector molecules. Typically, studies aiming to identify virulence factors such as effectors simulate a host interaction environment as plant pathogens primarily express these molecules at early stages of infection to overcome host defence systems [32, 49, 50]. This data also complies with the GO, PFAM and EC assignments, as the majority of functional annotations indicated core metabolic functions and therefore are unlikely to have virulence or infection related functions.

The significantly lower codon adaptation indices of the new genes compared to detected V1 genes can suggest a limited rate of protein translation, which implies that over time optimised transcriptional levels have a selective advantage for gene expression. This can also be influenced by repeat- induced point mutations which can have implication on codon frequencies, and ultimately CAIs [51]. Additionally, recent lateral gene transfers can result in altered codon frequencies as these involve acquiring genes that have codons optimised for different species [52]. We also observed a bias of CAIs in V1 annotations confirmed by mass spectrometry compared to the CAIs of all V1 genes. These experimentally confirmed genes had higher CAIs, which indicates that the highly abundant proteins sampled in this study are translated with high efficiency. The unusual codon usage of MT820649 was likely due to a sequencing error as there is no evidence of the presence of an intron at this locus (based on sequence homology with other P*hytophthora* species) and there is a stop codon 6 bp upstream. The remaining 28 sequences had normal codon usage.

## Conclusion

The data generated by shotgun LC-MS/MS confirmed 2,764 previously annotated gene models from the *P. cinnamomi* draft genome using high quality mass spectra from a diverse range of sub-proteome fractions. The spectral data suggested potential errors in gene calling, and using the spectral data, we were able to alter 60 genes by extending and merging exons, and identify 23 previously undescribed annotations in the *P. cinnamomi* genome. This demonstrates that the correlation between genes called by methods *in silico* are not always correlated to protein products, with evidence of annotation error rates of 2% of the detected proteome. This work demonstrates there are effective ways to use proteomics to correct boundary discrepancies and discover new genes. To our knowledge, this study presents the first use of spectral data to edit and manually annotate an oomycete pathogen. As more spectral data is accumulated, we expect there will be additional changes to the annotation including the discovery of more new genes.

## Acknowledgments

Funding was provided by Proteomics International. We thank Dr Paula Moolhuijzen, Johannes Debler and Darcy Jones for their technical assistance.

## Additional files

Additional file 1- format: gff. Title: CDS coordinates of edited genes, Description: Gene coordinates of manually edited V1 genes. Gene identification names include the original ID from https://mycocosm.jgi.doe.gov/Phyci1 as a reference.

Additional file 2- format: xlsx, Title: Master table of genes edited by extending features, Description: Summary of sequence, sequence features, peptide support, codon usage and functional assignment of edited genes generated by extending features

Additional file 3- format: xlsx, Title: Master table of genes edited by merging features, Description: Summary of sequence, sequence features, peptide support, codon usage and functional assignment of edited genes generated by merging features.

Additional file 4- format: xlsx, Title: Master table of new genes, Description: Summary of sequence, sequence features, peptide support, codon usage and functional assignment of new genes.

## List of abbreviations

GO: Gene Ontology
PFAM: Protein Families
1D SDS-PAGE: 1 Dimensional Sodium Dodecyl Suphate Polyachrilamide Gel Electrophoresis
RP-HPLC: Reverse phase High pressure liquid Chromatography
LC-MS: Liquid Chromatography-Mass Spectrometry
MS: Mass Spectrometry
TOF: Time of Flight
CONF: Confidence
FDR: False Discovery Rate
CDS: Coding Sequence
CAI: Codon Adaptation Index
GAPDH: Glyceraldehyde Phosphate Dehydrogenase
JGI: Joint Genome Institute
ORF: Open Reading Frame
KEGG: Kyoto Encyclopaedia of Genes and Genomes
KO: KEGG Orthologues
EC: Enzyme Code References

